# LMO2 regulates epithelial-mesenchymal plasticity of mammary epithelial cells

**DOI:** 10.1101/2025.05.22.655436

**Authors:** Veronica Haro-Acosta, Maria A. Juarez, Isobel J. Fetter, Andrew Olander, Shaheen S. Sikandar

## Abstract

Cellular plasticity in mammary epithelial cells enables dynamic cell state changes essential for normal development but can be hijacked by breast cancer cells to drive tumor progression. However, the molecular factors that maintain cellular plasticity through the regulation of a hybrid cell state (epithelial/mesenchymal) are not fully defined. As LMO2 has been previously shown to regulate metastasis, here we determined the role of LMO2 in the normal mammary epithelial cells. Using lineage tracing and knockout mouse models we find that *Lmo2* lineage-traced cells persist long-term in the mammary gland, both in the luminal and basal layer but have limited proliferative potential. *Lmo2* loss does not impact mammary gland development, but acute deletion decreases *in vivo* reconstitution. Moreover, LMO2 knockdown in mouse and human mammary epithelial cells (MECs) reduces organoid formation. We find that LMO2 maintains a hybrid cell state in MECs and LMO2 knockdown promotes mesenchymal differentiation. Transcriptional profiling of LMO*2* knockdown cells reveals significant enrichment in the epithelial-mesenchymal transition (EMT) pathway and upregulation of MCAM, a negative regulator of regenerative capacity in the mammary gland. Altogether, we show that LMO2 plays a role in maintaining cellular plasticity in MECs, adding insight into the normal differentiation programs hijacked by cancer cells to drive tumor progression.

## Introduction

Cellular plasticity has recently been recognized as a key hallmark in tumorigenesis and tumor progression^1, 2^. Epithelial-mesenchymal plasticity (EMP) is a form of cellular plasticity that allows cells to exist along a spectrum from epithelial to mesenchymal states, creating hybrid cellular states^1, 3, 4^. This state is now recognized as important for inducing stem-cell like properties, drug resistance and metastasis in cancer cells^2, 5–8^.

Mammary epithelial cells (MECs) have been used extensively as a model system to study regulators of cellular plasticity^9–11^. The mammary gland is a dynamic organ, with most of the development occurring postnatally, puberty and pregnancy, in response to hormones^12^. It is a bi-layered gland of luminal and basal epithelial cells where luminal layer gives rise to milk producing cells and basal cells have contractile function to enable the release of milk^12^. The CD49f^high^/EpCAM^med^ basal/myoepithelial cells in the mammary gland are enriched for *in vivo* reconstitution potential^13, 14^ and display characteristics of quasi-mesenchymal states (plastic cell state) with expression of both epithelial markers (EpCAM) and mesenchymal markers (smooth muscle actin)^15, 16^. The molecular regulation of the quasi-state in MECs remains poorly understood, although it has been an active area of research due to its potential implications in tumor development and progression.

We have previously demonstrated that the hematopoietic stem cell factor and transcriptional adaptor protein^17^, LMO2, is enriched in a minority population of immature tumor cells^8^. Our published results demonstrate that LMO2+ cells are metastatic, predict poor distant recurrence-free survival in patients and LMO2 is functionally required for metastasis^8^. However, the function of LMO2 in normal mammary epithelial cells is unknown. Here we explore the role of LMO2 in the normal mammary gland homeostasis using lineage tracing, knockout models, organoids assays and RNA-sequencing to find that LMO2 is not essential for mammary gland development but regulates cellular plasticity under regenerative conditions.

## Results

### LMO2 marks a minority long-lived population in the mammary gland

To determine cell fate and lineage hierarchy of *Lmo2* expressing cells in the mammary gland, we generated *Lmo2^CreERT^*^2^*/Rosa^mTmG^* mice (**Fig. 1A**). To test whether *Lmo2^CreERT^*^2^ marks *Lmo2* expressing cells we pulsed the mice at 8 weeks with a single dose of tamoxifen and sorted unswitched (TdTomato+) and switched (GFP+) cells. Gene expression profiling of the GFP*+* cells after a 36h pulse showed significant enrichment in expression of *Lmo2,* thus validating the *Lmo2 ^CreERT^*^2^ mouse model (**Fig. 1B**). Moreover, flow cytometry analysis of the mammary gland showed that *Lmo2*+ cells were present both in the luminal and basal lineages of the mammary epithelium (**Fig. 1C, Supplementary Fig. 1A, B**). To understand if *Lmo2*+ cells contributed to the developing mammary gland we pulsed the mice with tamoxifen at puberty (P28) followed by analysis at 12 weeks. Although we could detect *Lmo2-*lineage traced cells in the luminal and basal layers, we only observed small clusters of 2-3 cells (**Fig. 1D**). These data suggest that, while *Lmo2*+ cells are present during the development of the ductal epithelium, they have limited proliferative capacity (**Fig. 1D**). Similarly, pulsing the mice at 8-weeks and analysis at 16 weeks demonstrated a very small population of persistent *Lmo2*-lineage traced cells (**Fig. 1E**). To test whether the *Lmo2*+ cells expand during pregnancy, we pulsed female mice at 4 weeks, mated at 8 weeks, and analyzed at gestation day 15.5. We found that the *Lmo2-*lineage traced population expanded during pregnancy and were enriched in alveolar structures (**Fig. 1F**). However, in the second round of pregnancy no GFP+ cells were detected. Together, these data demonstrate that a minority population of *Lmo2*-expressing cells are present in both the basal and luminal lineages and *Lmo2*-lineage traced cells exist long-term in the mammary gland.

**Figure 1:**
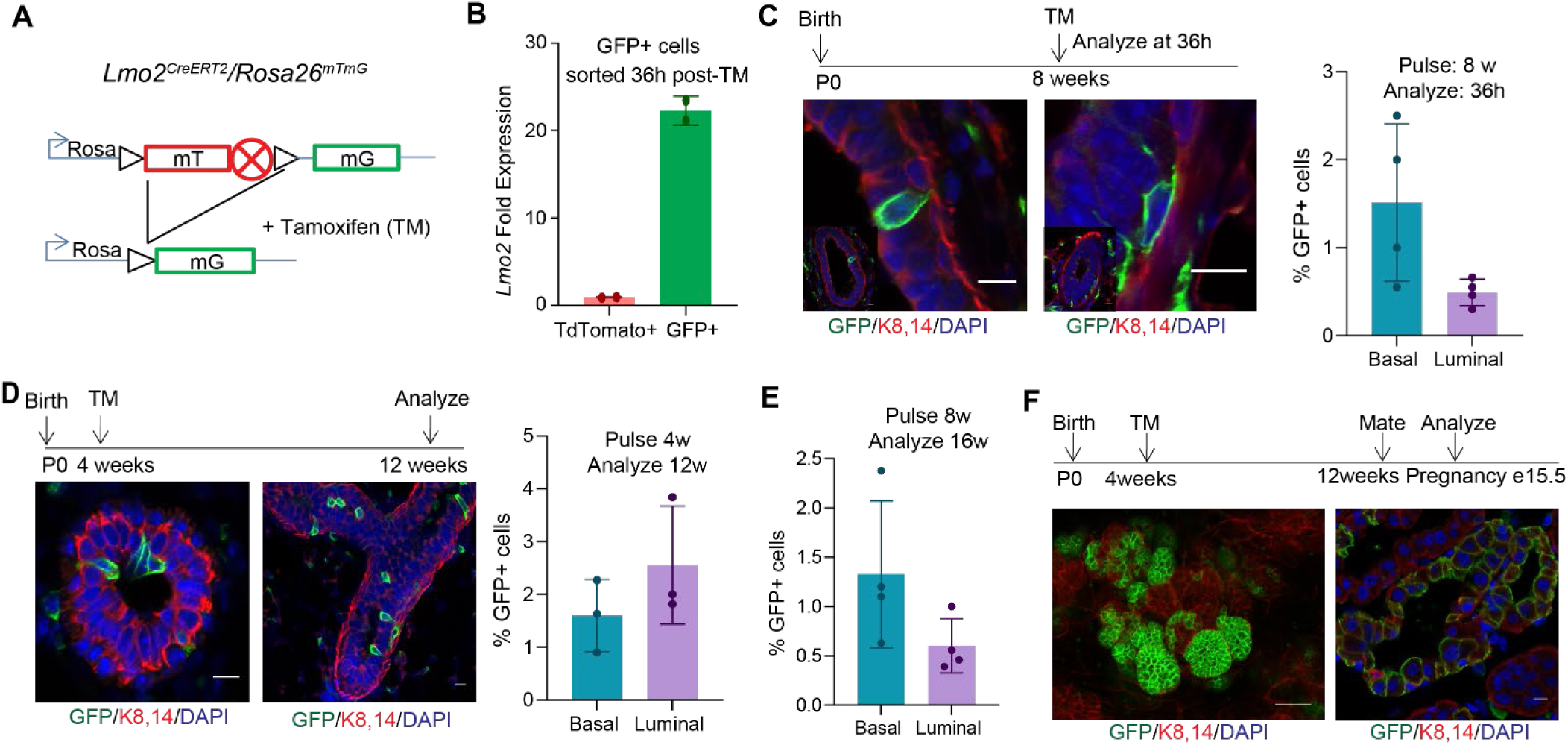
LMO2 marks a long-lived population in the mammary gland. (**A**) Schematic diagram showing the crossing of *Rosa26^mTmG^* reporter with *Lmo2^creERT2^* mice to generate *Lmo2^creERT2^*/ *Rosa26^mTmG^* (**B**) q-PCR analysis of GFP+ and GFP- cells sorted from *Lmo2-cre* mice (n=2) (**C**) Immunofluorescence analysis of adult *Lmo2^creERT2^*/ *Rosa26^mTmG^* mice pulsed with Tamoxifen at 8 weeks and analyzed after 36h. Representative images (left), quantification using flow cytometry (right) (n=4 mice) (see Supplementary Fig. 1B for flow cytometry plots). (**D**) *Lmo2^creERT2^*/ *Rosa26^mTmG^* pulsed at 4 weeks and analyzed at 12 weeks. Representative images (left), quantification by flow cytometry (right) (n=3 mice) (**E**) Flow cytometry quantification of GFP+ cells from *Lmo2^creERT2^*/ *Rosa26^mTmG^* mice pulsed at 8 weeks and analyzed at 16 weeks (n=4 mice). (**F**) Representative images of *Lmo2^creERT2^*/ *Rosa26^mTmG^* mice pulsed at puberty, mated 8 weeks later and analyzed at gestation day 15.5.

### LMO2 regulates *in vivo* reconstitution potential and organoid formation of MECs

We next sought to determine whether LMO2 is required for normal mammary gland development. Germline *Lmo2* deletion in mice causes embryonic lethality due to failure in fetal hematopoiesis^18^. Hence, to understand if LMO2 is necessary for the formation of the ductal epithelium, we crossed *Lmo2^fl/fl^* mice^19^ with *Keratin14^Cre^* mice^20^ to specifically delete *Lmo2* in the epithelial compartment. *Keratin14^Cre^/Lmo2^fl/fl^*mice developed a normal epithelial tree with a mild developmental delay at 5 weeks that was subsequently rescued by week 8 (**Supplementary Fig. 2A**). FACS analysis of the gland showed no difference in the ratio of basal to luminal cells in *Keratin14^Cre^/Lmo2^+/+^* and *Keratin14^Cre^/Lmo2^fl/fl^* in 8–10 week old mice (**Fig. 2A, B**).

**Figure 2:**
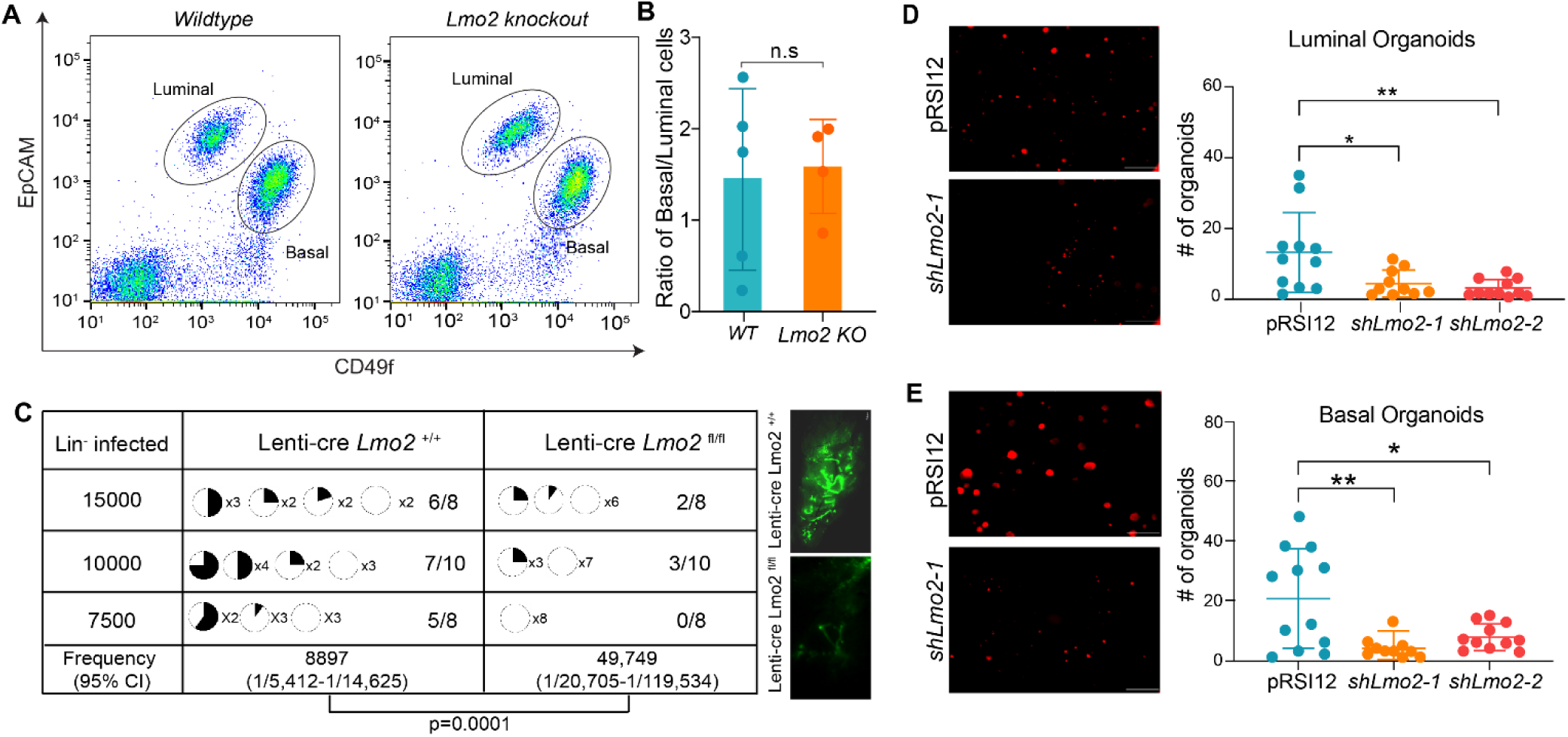
*Lmo2* loss does not impact normal development but leads to decreased reconstitution potential. (**A**) Flow cytometry analysis of Live/Lineage^-^ cells from *K14^Cre^/Lmo2^fl/fl^* and *K14^Cre^/Lmo2^+/+^* mammary glands (see Supplementary Fig. 1A for gating strategy). Representative flow plot of Live/Lineage-cells. (**B**) Ratio of basal to luminal cells in the mammary epithelial cell population from *K14^Cre^/Lmo2^fl/fl^*and *K14^Cre^/Lmo2^+/+^* mammary glands (n=5 mice) (**C**) Limiting dilution transplantation assay with lineage depleted cells from *Lmo2^+/+^* and *Lmo2^fl/f^*^l^ mice transduced with Lenti-cre-GFP plasmid. Pie chart indicates percentage of fat-pad filled. The proportion of positive outgrowths out of the total cell number injected is shown on the right. (Note: Injections with 5000 or less lineage depleted MECs transduced with lenti-cre did not lead to any outgrowths in either *Lmo2^+/+^* or *Lmo2^fl/fl^* mice.) P-value and repopulating cell frequency was calculated using ELDA. Representative images of transplantation outgrowth are displayed on the right. (**D**) Organoid formation in control and *Lmo2* knockdown basal epithelial cells sorted from 3-month-old mice. Representative image (left), quantification (right), n=3 mice with 3 technical replicates each. (**E**) Organoid formation in control and *Lmo2* knockdown luminal epithelial cells sorted from 3-month old mice. Representative image (left), quantification (right), n=3 mice with 3 technical replicates each. Data are shown as mean ± SD. Statistical significance was calculated using unpaired student’s t test (B) and ordinary one-way ANOVA with multiple comparison test (D). * p < 0.05, ** p < 0.01, *** p < 0.001, **** p<0.0001.

To test whether *Keratin14^Cre^* leads to efficient deletion of *Lmo2*, we performed genotyping and q-PCR analysis. While we could detect the deletion product (**Supplementary Fig. 2B**), q-PCR analysis on sorted MECs showed that the *Lmo2* transcript is expressed in MECs sorted from *Keratin14^Cre^/Lmo2^fl/fl^*mice, albeit at low levels (**Supplementary Fig. 2C**). This could be due to an inherent inefficiency of Cre that allows cells to escape *Krt14*-Cre-mediated deletion and prevents us from fully assessing the impact of *Lmo2* deletion in the mammary gland.

Lineage tracing measures a cell’s fate under normal homeostasis while reconstitution assays measure the potential of the cell under austere regenerative conditions^21, 22^. The ability of CD49f^high^/EpCAM^med-low^ basal cells to regenerate the mammary gland upon transplantation^13, 14^ reflects an intrinsic plasticity of the cells. To test whether *Lmo2* loss alters reconstitution potential, we used a GFP-tagged lentiviral-cre-mediated deletion in MECs and performed transplantation assays. We found that acute loss of LMO2 significantly reduced *in vivo* reconstitution potential, leading to both fewer and less filled mammary fat pads, suggesting that *Lmo2* contributes to regeneration of the mammary gland *in vivo* (**Fig. 2C**).

To confirm our findings using organoid models, we also generated shRNA against *Lmo2* (**Supplementary Fig. 2D**) and transduced sorted luminal and basal cells with either a control vector or shRNA against *Lmo2*. Consistent with our *in vivo* transplantation data, we find that *Lmo2* knockdown significantly reduced organoid forming capacity of both luminal (**Fig. 2D**) and basal cells (**Fig. 2E**). Taken together, these data implicate LMO2 as a positive regulator of *in vivo* regeneration and clonogenic potential in mammary epithelial cells.

### LMO2 maintains a hybrid cell state in mammary epithelial cells

To test whether LMO2 has a role in normal human mammary epithelial cells, we used the MCF10A cell line, that have been extensively used to study breast tumor initiation and cellular plasticity as they consist of a heterogeneous population of cells^23, 24^. LMO2 knockdown (**Supplementary Fig. 3A**) significantly reduced MCF10A colony formation in 2D (**Fig. 3A**). Similarly, LMO2 knockdown significantly impaired acini formation in 3D culture (**Fig. 3B**), suggesting a role for LMO2 in supporting regenerative growth under clonogenic conditions in human mammary epithelial cells.

**Figure 3:**
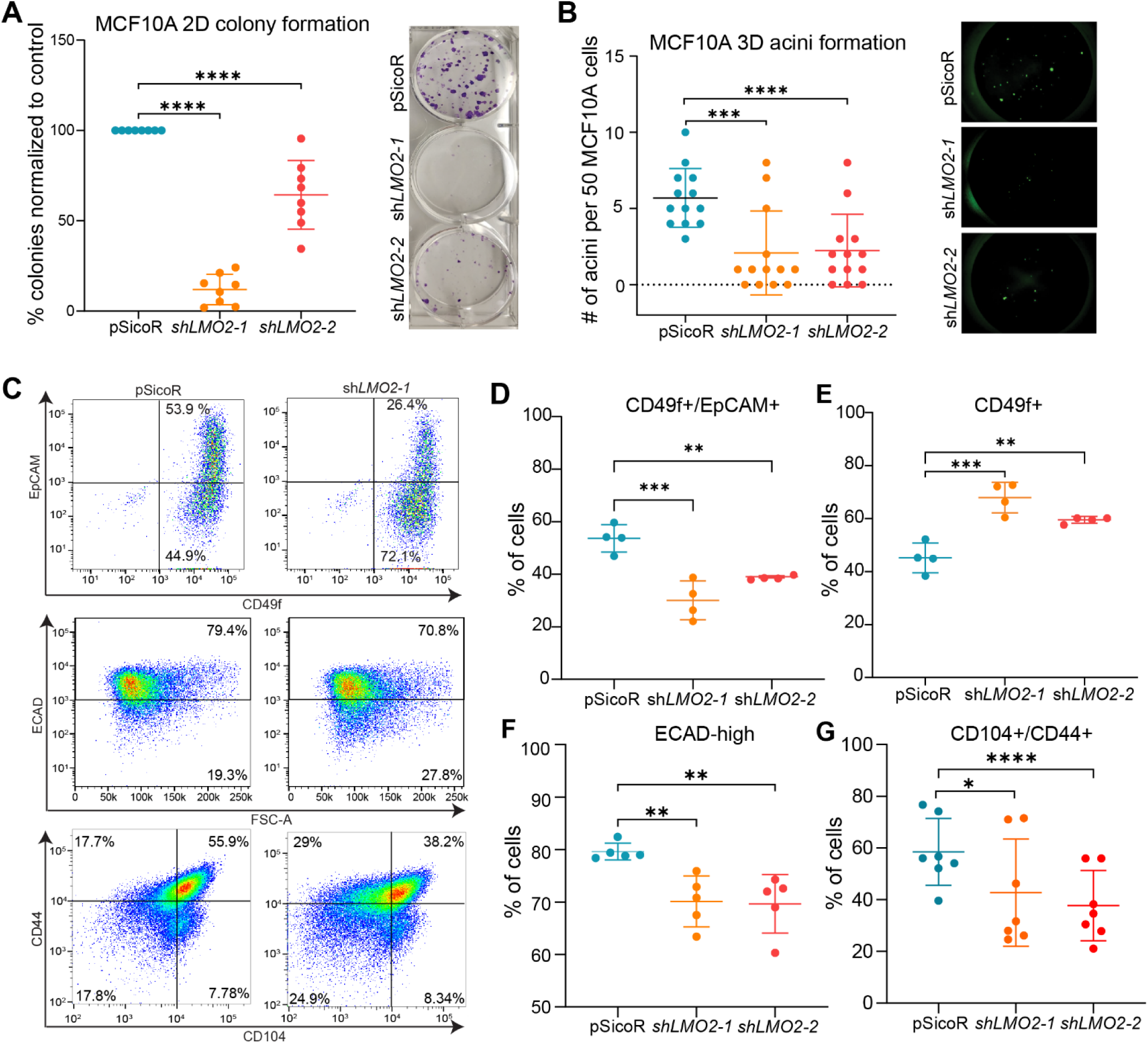
*LMO2* knockdown in MCF10A cells reduces organoid formation, hybrid cell state and induces mesenchymal differentiation. **A**) Quantification (left) and representative images (right) of colony formation in control and *LMO2* knockdowns MCF10A cells (n=3 biological and 3 technical replicates). (**B**) Quantification (left) and representative images (right) of acini formation of control and *LMO2* knockdown MCF10A cells. (n=3 biological and 3 technical replicates). (**C**) Representative images of flow cytometry analysis of control and *LMO2* knockdown MCF10A cells with previously established markers EpCAM and CD49f (top row), ECAD (middle row), CD44 and CD104 (bottom row). (**D-G**) Quantification of flow cytometry analysis of control and *LMO2* knockdown MCF10A cells for the following populations: (**D**) EpCAM+/CD49f+ (**E**) CD49f+ (**F**) ECAD-high (**G**) CD104+/CD44+. Data are shown as mean ± SD. Statistical significance was calculated using ordinary one-way ANOVA with multiple comparison test. * p < 0.05, ** p < 0.01, *** p < 0.001, **** p<0.0001.

To test whether decrease in colony and acini formation with LMO2 knockdown is due to changes in proliferative potential, we performed WST-1 assays. LMO2 knockdown in MCF10A cells had no effect on proliferation (**Supplementary Fig. 3B**). To understand whether LMO2 loss induces apoptosis or decreases cell viability, we performed flow cytometry analysis on MCF10As at day 4, 7 and 11 post-transduction with either a control vector or *LMO2* knockdown. We found no significant differences in apoptosis by Annexin V staining or cell viability at all the different time points examined (**Supplementary Fig. 3C**). Thus, our data suggest that LMO2 promotes clonogenicity through mechanisms independent of proliferation or survival pathways.

Recent studies have shown that cells expressing both epithelial and mesenchymal markers (hybrid, E/M) have higher regenerative and clonogenic capacity than their differentiated counterparts^5, 25–29^. To test whether LMO2 knockdown impacts the hybrid cell state, we performed flow cytometry analysis with markers previously used to identify epithelial and mesenchymal/basal cells, CD49f and EpCAM, respectively^16^. We found that MCF10A cells primarily consist of two populations: a double positive population (EpCAM+/CD49f+) and a basal/mesenchymal population (EpCAM-/CD49f+). Interestingly, LMO2 knockdown significantly reduced the percentage of cells that are EpCAM+/CD49f+ and increased the percentage of CD49f+ cells (**Fig. 3C top panel, D, E**), suggesting differentiation towards a basal/mesenchymal state. Similarly, we used E-cadherin as a marker for epithelial cells and found that LMO2 knockdown reduced the percentage of E-cadherin+ cells (**Fig. 3C middle panel, F**). To further understand whether LMO2 regulates the intermediate epithelial-mesenchymal state, we utilized CD104 and CD44, which have been previously demonstrated to mark hybrid E/M cells in breast cancer^30^. We found that LMO2 knockdown reduced the proportion of cells in the hybrid state as indicated by decrease in CD104+/CD44+ cells (**Fig. 3C bottom panel, G**). Collectively, our data suggests that LMO2 regulates a hybrid cell state in MCF10A cells and LMO2 loss promotes differentiation towards the basal/mesenchymal state.

### RNA-sequencing identifies molecular pathways regulated by LMO2

As LMO2 is a transcriptional adaptor protein, we next sought to identify transcriptional programs regulated by LMO2 that maintain a hybrid cell state in MCF10A cells. Hence, we performed RNA-sequencing analysis on control and LMO2 knockdown cells at 6-days post-transduction (**Fig. 4A**). We found a significant increase in the expression of genes associated with mesenchymal differentiation such as *FN1, MCAM,* and *FBN1* and down regulation of genes involved in maintaining stem/progenitor cells such as *ALHD1A3, LIF* and *CSF* (**Fig. 4B**). MCAM has been demonstrated as a negative regulator of clonogenicity and regenerative capacity of mammary epithelial cells, where MCAM loss promotes recruitment of macrophages and expansion of MECs through non-canonical Wnt signaling^31^. To validate RNA-sequencing analysis we performed flow cytometry for MCAM and found that LMO2 knockdown increases the percentage of MCAM+ MCF10A cells (**Fig. 4D, E**), although it did not reach statistical significance for the second shRNA construct.

**Figure 4:**
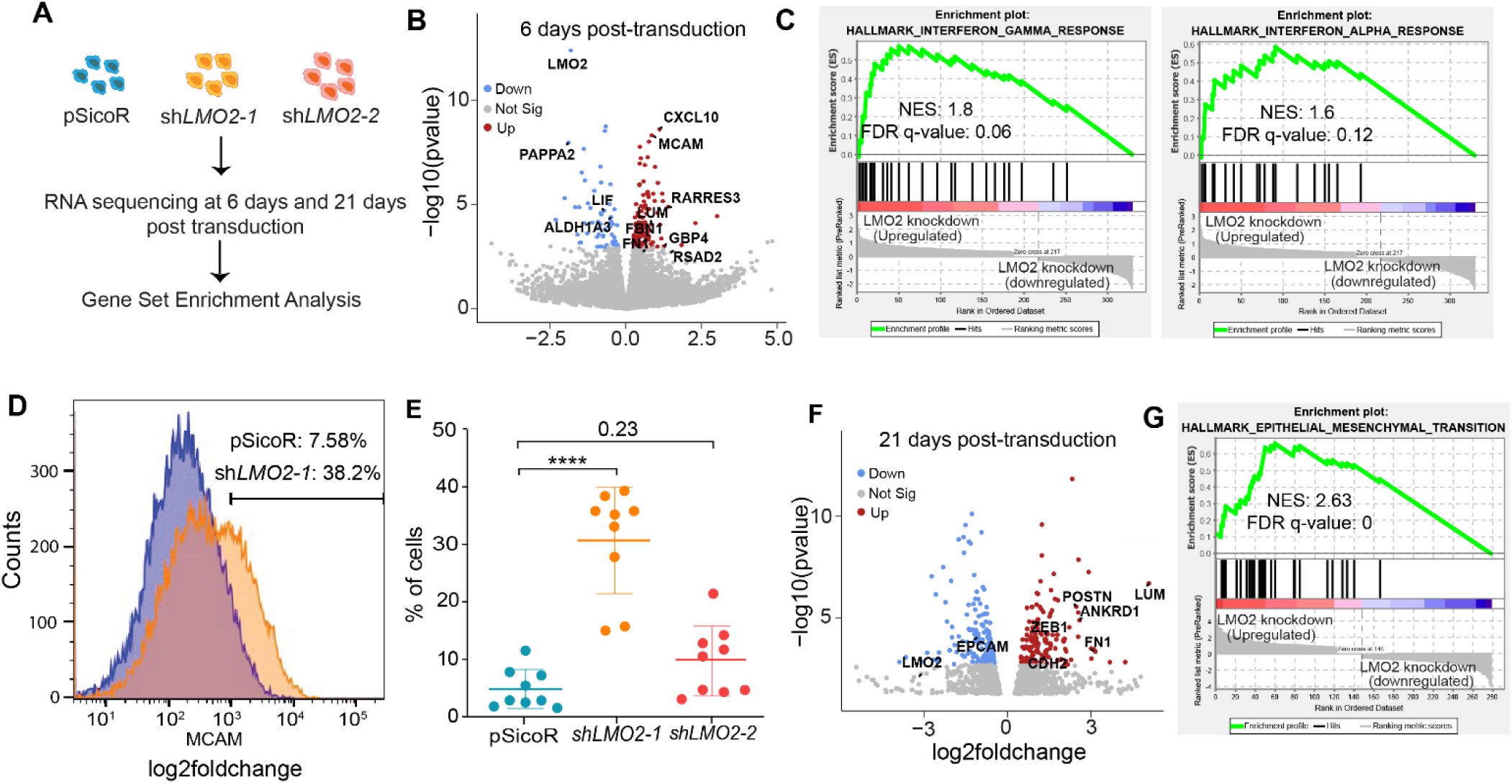
Transcriptomic analysis of *LMO2* knockdown in MCF10A cells. (**A**) Schematic of bulk-RNA sequencing of control and *LMO2* knockdown MCF10A cells (n=3) at 6 and 21 days after infection. (**B**) Volcano plots visualizing the fold-changes and p-values between control and *LMO2* knockdown after 6 days post-transduction. (**C**) Enrichment plots for interferon-gamma and interferon-alpha response gene sets from data displayed in **B** at 6 days post-transduction. (**D**) Representative flow cytometry analysis of MCAM expression in control and *LMO2* knockdown MCF10A cells. (**E**) Quantification of flow cytometry analysis of MCAM expression in control and *LMO2* knockdown MCF10 cells (n=9). (**F**) Volcano plots visualizing the p-value and fold change between control and *LMO2* knockdown MCF10A cells 21 days after post-transduction (**G**) Enrichment plot of the epithelial-mesenchymal transition gene set from data displayed in **F** at 21 days post-transduction. NES = normalized enrichment score, FDR = false discovery rate. Statistical significance was calculated using ordinary one-way ANOVA with multiple comparison test. * p < 0.05, ** p < 0.01, *** p < 0.001, **** p<0.0001.

Moreover, we saw an increase in expression of genes involved in antigen presentation (*HLA-A, HLA-B* and *HLA-F*) and interferon response (*CXCL10, RSAD2,* **Fig. 4B, Supplementary Table 1**). Gene set enrichment analysis (GSEA)^32^ using the hallmark geneset showed an upregulation of pathways associated with interferon alpha and gamma signaling (FDR q-value <0.25) (**Fig. 4C, Supplementary Table 1**), potentially reflecting transcriptional programs that are necessary for hematopoietic stem cell maintenance^33–35^. Although there was an upregulation of specific mesenchymal markers, epithelial to mesenchymal transition was not significantly enriched in the GSEA (FDR q-value = 0.28) (**Supplementary Table 1**).

The transition of epithelial-mesenchymal transition occurs over time^36–38^. To further characterize this phenotypic shift, we performed RNA-sequencing analysis at 21 days post-transduction with the first shRNA construct. Consistent with our flowcytometry analysis, we found a significant upregulation of mesenchymal genes such as *ZEB1, FN1, LUM, CDH2* and *POSTN*^39^ (**Fig. 4F**). Moreover, the only significantly enriched Hallmark geneset in *LMO2* knockdown was epithelial to mesenchymal transition (**Fig. 4G, Supplementary Table 1**). Thus, our data demonstrates that *LMO2* knockdown induces a mesenchymal cell state and promotes upregulation of *MCAM*, a negative regulator of MEC regenerative capacity.

## Discussion

Cellular plasticity is key to maintaining regenerative capacity in the mammary gland and plays an important role in tumor progression^2, 9–11, 40^. Previous studies have implicated several important transcriptional regulators of EMT as critical regulators of this process, such as Snail, Slug, Twist1 and Zeb1^41–43^. However, the role of other proteins beyond the core EMT transcription factors are only beginning to be understood.

Our previous studies have shown that LMO2 marks a population of immature tumor cells and functionally LMO2 is required in the early steps of the metastatic process^8^. The role of LMO2 in the normal mammary gland, identified here, aligns with the known role of LMO2 in promoting metastasis rather than primary tumor growth. Notably, the *in vivo* transplantation assay recapitulates key features of the metastatic process, as successful engraftment requires activation of programs involved in invasion, survival, and proliferation within a foreign microenvironment. Our data is also consistent with previous studies showing that deletion of Snai2 or Twist1 in mammary epithelial cells does not significantly impact formation of the ductal tree *in vivo* but impacts *in vivo* reconstitution^44–48^. While we find that *Lmo2* knockout does not impact normal mammary gland development, we are unable to unequivocally measure the full impact of *Lmo2* deletion *in vivo* given that there might be some cells that do not undergo recombination.

Epithelial-to-mesenchymal plasticity and maintenance of a hybrid cell state has been recently shown to be a critical factor for tumorigenicity, chemoresistance and metastatic potential in cancer cells^6, 7, 29, 49–52^. Since tumor progression often co-opts developmental and regenerative processes, elucidating these processes under normal conditions is crucial for identifying how they are hijacked during tumorigenesis. Here, we find that loss of LMO2 in mammary epithelial cells pushes cells towards a mesenchymal differentiated state that is less regenerative^5, 13, 14, 29^. LMO2 loss induces expression of several mesenchymal markers, including MCAM. Interestingly, loss of MCAM has the opposite effect where it increases clonogenicity and regenerative capacity of MECs, through the IL4-Stat6 axis and non-canonical Wnt signaling^31^. Although our study does not determine the precise mechanism of LMO2 in MECs, we speculate that LMO2 promotes cellular plasticity in part through suppression of *MCAM* expression. Collectively, our study reveals a previously unknown role for LMO2 in maintaining regenerative capacity of mammary epithelial cells and a hybrid cell state.

## Materials and Methods

### Mice

All mice used for this study were maintained at the UCSC Animal Facility/Vivarium in accordance with the guidelines of the Institutional Animal Care and Use Committee (Protocol #Sikas2311dn). *Lmo2^CreERT2^* and *Lmo2^fl/fl^*mice were a gift from Dr. Terence Rabbits, Tg(KRT14-cre)1Amc/J (RRID:IMSR_JAX:004782, Strain #018964) and (ROSA)26Sor^tm4(ACTB-tdTomato,-EGFP)Luo^/J (RRID:IMSR_JAX:007676) were purchased from JAX labs. *Lmo2*^CreERT2^/*Rosa26*^mTmG^ BL6 female mice were subjected to intraperitoneal injection of Tamoxifen. Tamoxifen was dissolved in corn oil at a concentration of 10 mg/ml. Each mouse received a dose of 1.5 mg, as previously described^8^. Mice were injected at 4 or 8 weeks and subsequently analyzed at 36 hours, 12 weeks, or 14 weeks post-injections.

The mammary glands were either stained and imaged or enzymatically digested and subjected to flow cytometry. Krt14-cre mice were crossed to *Lmo2^fl/fl^* to generate mammary specific deletion of Lmo2.

### Cell Lines

MCF10A cells were maintained in Advance DMEM 1X/F12 media (Fisher Scientific: # 12634010) with 5% of Horse serum (Life Technologies: # 16050-122), 1% of PSA (Fisher Scientific: ICN1674049), 100ug/ml of human EGF (Peproptech: # AF-100-15-100UG), 100 ng/ml of Cholera toxin (Sigma Aldrich: # C8052-0.5MG), 9.5-11 mg/ml of Insulin (Neta Scientific: # SIAL-I9278-5ML), 5ug/ml of Hydrocortisone (Sigma-Aldrich Company: # H0888-1G). HEK293T cells were maintained with DMEM at 10% Fetal bovine serum (Fisher Scientific: # MT350110CV) and 1% PSA (Fisher Scientific: ICN1674049). Cultures were maintained at 37°C with an atmosphere of 5% CO2. Images of cells were taken on a Zeiss Live Microscope capturing both brightfield and fluorescence.

### Mammary Gland Digestion

L2-5 and R2-5 mammary glands were harvested, minced, and chemically digested overnight in Advanced DMEM F/12 with 1% PSA, gentle collagenase/hyaluronidase (Stem Cell Technologies: 7919), and DNAse I at 37°C, 5% CO2, and added humidity as previously described72. Briefly, partially digested glands were then mechanically digested by pipetting with a serological pipette until no tissue pieces were visible. Digested glands were washed with staining buffer (Hank’s Balanced Salt Solution, 2% Bovine Calf Serum, 1% PSA) and centrifuged (1500RPM) at 4°C for 5 minutes. Red blood cells were lysed with 5mL of ACK Lysis buffer for 5 minutes, and cells were washed with 15mL of staining buffer. Cells were treated with 0.25% Trypsin with EDTA and gently pipetted continuously for 2-3 minutes to digest the basement membrane. Cells were then treated with DNAse I and Dispase and pipetted continuously for 2-3 minutes to prevent clumping. The single-cell suspension was then filtered through a 40μm mesh strainer and pelleted via centrifugation (1500RPM, 4°C, 5 minutes). Cells were then resuspended in staining buffer and transferred to FACS tubes for staining.

### Transplantation Assay

Live lineage^neg^ cells were sorted from *Lmo2^+/+^*and *Lmo2^fl/fl^* mice. Gating strategy as shown in Supplementary Fig 1. Cells were transduced with Lenti-cre-GFP overnight and then counted before transplantation. Cells were resuspended with 33% Matrigel injected into the cleared mammary fat pad at 15,000, 10,000, and 7,500. Per transplant 10μL was injected into cleared fat pads of weaning age C57BL/6 mice (21–28 days) as previously described^29^. All transplants were allowed to grow 8 weeks before analysis.

### Flow Cytometry Analysis

Cells were stained with antibodies listed in **Supplementary Table 2** for 15 minutes at room temperature, as previously described^29^. Stained cells were then washed with a staining buffer, resuspended with DAPI (1:10,000), filtered, and analyzed on a BD Biosciences FACSAria cell sorter. See **Supplementary Fig. 1A** for FACS gating strategies. MCF10A infected with pSicoR, sh*LMO2-1*, and sh*LMO2*-2 were resuspended in staining buffer consisting of HBSS (Fisher Scientific: # MT21022CV) with 2% BCS (Sigma Aldrich: # 12133C-500ML) and 1% PSA (Fisher Scientific: ICN1674049) and stained with antibodies shown in **Supplementary Table 2**. Stained cells were then washed with staining buffer, resuspended with DAPI (1:10,000), filtered, and analyzed on a BD Biosciences FACSAria cell sorter. Data were analyzed using FlowJo software (10.10.0).

### Mammary Organoids

Luminal and basal cells were sorted from the mammary gland of 3 month old C57/Bl6 mice as shown in **Supplementary Fig 1A**. 96-well low attachment plate (Fisher Scientific: 7200603) plates were used to seed a 50μL of Growth Factor Reduced Matrigel (Fisher Scientific: # CB40230C) mixed with 11,000 L-Wnt3a irradiated cells per well. Luminal or basal cells were transduced with pRSI12, sh*Lmo2-2*, or sh*Lmo2-1* and seeded at 2500 per well. Organoid media was replenished every two days. The organoid media consisted of Advanced DMEM 1X/F12 media (Fisher Scientific: # 12634010), 1% PSA (Fisher Scientific: ICN1674049), 10% FBS (Fisher Scientific: # MT350110CV), 50 ng/ml human EGF (Peprotech: # AF-100-15-100UG),100 ng/ml mouse Noggin (Peprotech: # 10773-428), 250 ng/ml Human R-spondin-1 (Peprotech:120-38-50UG), 100x N-2 supplement (Life Technologies: # 17502048), 50x B-27 supplement (Life Technologies: # 17504044), 10mM HEPES (Fisher Scientific: # 12634028), 10uM Y-27632 (Fisher Scientific: # 125410), 1x GlutaMAX (Life Technologies: # 35050061). Organoids were maintained at 37°C with an atmosphere of 5% CO2. Images of cells were taken on a Zeiss Live Microscope capturing fluorescence. The images were uploaded into Biodock, and the AI was trained to quantify the number and size of the organoids.

### Collagen/Matrigel Acini formation

The protocol was modified from Brugge Lab (https://brugge.hms.harvard.edu/protocols). 250μL of Collage I (Fisher Scientific: #354249) was neutralized by adding 31μL of sterile 10x PBS and sterile NaOH until Collagen I had a pH of 7.5. Growth Factor Reduced Matrigel (Fisher Scientific: # CB40230C) was mixed with neutralized Collagen I at 4:1 and 40μl was seeded in a 96-well low attachment plate. MCF10A cells were infected with lentiviruses containing pSicoR, sh*LMO2-1*, or sh*LMO2-2* and seeded in MCF10A growth media with 5% horse serum (Life Technologies: # 16050-122) at 2% Growth Factor Reduce Matrigel and hEGF (5ng/mL, Peprotech: # AF-100-15-100UG) at a density of 50 cells/well. The acini were imaged on a Keyence BZ-9000 microscope The images were uploaded into Biodock, and the AI was trained to quantify the number and size of the organoids.

### Library Construction, Quality Control and Bulk RNA Sequencing

Bulk RNA sequencing was performed by Novogene on MCF10A cells transduced with either control (pSicoR) or shLMO2 knockdown vectors. RNA was isolated according to manufacturer’s instructions (Qiagen RNEasy Plus Micro Kit, Cat. No. 74034). Messenger RNA was purified from total RNA using poly-T oligo-attached magnetic beads. After fragmentation, the first strand cDNA was synthesized using random hexamer primers, followed by the second strand cDNA synthesis using either dUTP for directional library or dTTP for non-directional library. For the non-directional library, it was ready after end repair, A-tailing, adapter ligation, size selection, amplification, and purification. For the directional library, it was ready after end repair, A-tailing, adapter ligation, size selection, USER enzyme digestion, amplification, and purification. The library was checked with Qubit and real-time PCR for quantification and bioanalyzer for size distribution detection. Quantified libraries will be pooled and sequenced on Illumina platforms, according to effective library concentration and data amount.

### Data Quality Control

Raw data (raw reads) of fastq format were processed through fastp software. In this step, clean data (clean reads) were obtained by removing reads containing adapter, reads containing ploy-N and low quality reads from raw data. Q20, Q30 and GC content were calculated. All the downstream analyses were based on clean data with high quality.

### Reads mapping to the reference genome

Reference genome (GRCm39/mm39) and gene model annotation files were downloaded from genome website directly. Index of the reference genome was built using Hisat2 v2.0.5 and paired-end clean 1 reads were aligned to the reference genome using Hisat2 v2.0.5. We selected Hisat278 as the mapping tool for which Hisat2 can generate a database of splice junctions based on the gene model annotation file and thus a better mapping result than other non-splice mapping tools.

### Quantification of gene expression level

FeatureCounts79 v1.5.0-p3 was used to count the reads numbers mapped to each gene. Gene expression was then converted to transcripts per million (TPM) by normalizing for gene length, then sequencing depth. TPM was chosen as a normalization method as it allows for the more accurate comparison across samples.

### Differential expression analysis and gene set enrichment analysis

Differential expression analysis of two conditions (three biological replicates per condition) was performed using the DESeq2Rpackage81 (1.20.0). DESeq2 provides statistical routines for determining differential expression in digital gene expression data using a model based on the negative binomial distribution. Differentially expressed genes were identified by comparing control and *LMO2* knockdown cells. The resulting P-values were adjusted using Benjamini-Hochberg’s approach for controlling the false discovery rate. Genes with an adjusted P-value <=0.1 found by DESeq2 were assigned as differentially expressed. Heatmaps. GSEA was performed on all DEGs at day 6 post-transduction and a preranked list of genes at 21 days post-transduction. The FDR q value < 0.25 was considered significant between control and knockdown conditions using the Broad Institute’s software.

### shRNA cloning and Virus production

The lentivirus plasmid pSicoR (Plasmid #11579, Addgene) was utilized as the backbone for generating lentivirus plasmids containing the human short hairpin RNA insertions targeting *LMO2*.The sequences for the LMO2 shRNA are 5′-GACGCATTTCGGTTGAGAA-3′ and 5′-GCATCCTGTGACAAGCGGATT-3′.

The lentivirus plasmid pRSI12 (Cellecta) was utilized for the backbone for generating lentivirus plasmids containing mouse short hairpin insertion targeting *Lmo2*. The sequences for mouse *shLmo2* 5’- TAATCTCCTAAGAAATGCCTC-3’ and 5’-AATTGCACAACTCTAGTCCAT-3’. HEK293T cells were seeded at a density of 5x10^6^ cells per 10cm plate the day before transfection. For lentivirus production, two separate cocktails were prepared. The first cocktail contained 15μg of lentivirus plasmid per plate, 7.5μg of pCMV-R8.91 per plate, and 5μg of pCMV-VSV-G per plate, diluted in 1.5ml of Opti-MEM reduced serum media (Life Technologies Corporations: #31985070). The second cocktail consisted of 1.5mL of Opti-MEM reduced serum media supplemented with 40μl of Lipofectamine 2000 (Fisher Scientific: #11668019) per plate. After a 5 minute incubation period, the two cocktails were mixed and incubated for 25 minutes before being added to the HEK293T cells. The transfection mixture was added to the HEK293T cells and incubated for 6 hours. Following incubation, the media was replaced with fresh growth media. After 48 hours, the lentivirus-containing media was collected and precipitated using Lentivirus Precipitate Solution (ALSTEM: #VC100). The precipitated lentivirus particles were then resuspended in an appropriate buffer for subsequent use in experiments or storage at -80°C.

### Crystal Violet Blue Colony Assay

MCF10A cells infected with lentivirus were seeded in a 6-well cell culture-treated plate at 500 cells per well. The cells were incubated at 37°C for two weeks. The cells were fixed with 10% neutral-buffered formalin for 15 minutes, then stained with 0.01% crystal violet for 60 minutes. The cells were rinsed with 1X PBS (Life Technologies: # 14190144), air dried, and the number of colonies counted.

### Annexin V Staining

MCF10A cells were transduced with pSicoR, sh*LMO2-1*, or sh*LMO2-2*and analyzed on days 4, 6, and 11. Culture media was removed, and cells were detached using 0.25% trypsin for 5 minutes at 37°C, then neutralized with MCF10A culture media. Cells were centrifuged at 1200 rpm for 5 minutes at 4°C, the supernatant was discarded, and the cell pellet was resuspended in 1X binding buffer (Invitrogen: # BDB556454) at a concentration of 1mL per 1 × 10⁶ cells. APC Annexin V antibody (BioLegend: #640920) was added at 5µL per 100µL of 1X binding buffer. Cells were incubated with the antibody for 15 minutes, followed by a wash with 3mL of 1X binding buffer. Cells were again centrifuged at 1200 rpm for 5 minutes at 4°C. The final cell pellet was resuspended in 1X binding buffer containing DAPI at a 1:10,000 dilution and analyzed by flowcytometry on the BD FACSAria.

### Whole Mount

Mammary glands were dissected and placed on Superfrost slides (Fisher Scientific: # 48382-200 (PK)) and placed in Carnoy’s fixative solution overnight. The following day, the slides were sequentially immersed in 70% ethanol and 50% ethanol for 15 minutes each. Afterward, they were rinsed in deionized water for 15 minutes and stained overnight with Carmine Alum. The next day, the slides were washed sequentially in 70%, 95%, and 100% ethanol for 15 minutes each. The glands were then cleared for 5 minutes to 1 hour, mounted on slides using Permount (Fisher Scientific: # SP15100) and allowed to dry overnight before imaging.

### WST-1 Assay

MCF10A cells were seeded at a density of 30,000 cells per well in a 12-well culture plate and transduced with either lentiviral constructs pSicoR, sh*LMO2-1*, or sh*LMO2-2*. Following transduction, cells were re-seeded at 1,000 cells per well in a 96-well plate for viability analysis. Cell viability was assessed every other day using the WST-1 reagent (Neta Science: # SIAL-5015944001). After a 2-hour incubation with WST-1, absorbance was measured at 440 nm using a microplate reader.

### Quantitative real-time PCR

Cells for qPCR were collected from sorted MECs or 2D tissue culture and spun down. RNA was extracted using the RNEasy Micro Kit (Qiagen: #74104) and quantified via ultraviolet spectrophotometer (Nanodrop). cDNA was synthesized using iScript cDNA Synthesis Kit (BioRad: #1708891), then underwent real-time polymerase chain reaction with SYBR Green Universal Master Mix (Fisher Scientific: # 4309155) for specific target genes. Primers used were *LMO2* F:5’-GGCCATCGAAAGGAAGAGCC-3’ and R:5’-GGCCCAGTTTGTAGTAGAGGC-3’, *Lmo2* F:5’-TCGCTCTCTCTCTTTGGCGT-3’ and R:5’- TGGCTTTCAGGAAGTAGCGG-3’, *ACTB*F:5’-CACCATTGGCAATGAGCGGTTC-3’ and R:5’- AGGTCTTTGCGGATGTCCACGT-3’, *GAPDH* F:5’-GTCTCCTCTGACTTCAACAGCG-3’ and R:5’-ACCACCCTGTTGCTGTAGCCAA-3’, *Actb* F:5’-GGCTGGATTCCCCTCCATCG -3’ and R:5’- CCAGTTGGTAACAATGCCATGT-3’, *Gapdh* F:5’-TGGCCTTCCGTGTTCCTAC -3’ and R:5’- GAGTTGCTGTTGAAGTCGCA -3’. All expression data was normalized to housekeeping controls.

### Statistical analysis

All graphs display the average as central values, and error bars indicate ± SD unless otherwise indicated. P values are calculated using paired or unpaired t test, ANOVA, Wilcoxon rank-sum test, and Mann-Whitney U test, as indicated in the figure legends. All P and Q values were calculated using Prism (10.2.2), unless otherwise stated. For animal studies, sample size was not predetermined to ensure adequate power to detect a prespecified effect size, no animals were excluded from analyses, experiments were not randomized, and investigators were not blinded to group allocation during experiments.

## DATA AVAILABILITY

Data generated or analyzed during this study are included in this published article (and its supplemental information files). Data needed to evaluate the conclusions in the paper are present in the paper and/or the Supplemental Materials. Source data files will be made available upon request. Bulk RNA-sequencing data generated in this study have been deposited in the Gene Expression Omnibus. All bioinformatics tools used in this study are published and publicly available.

## ACKNOWLEDGMENTS

We thank Bari Nazario and Patricia Lovelace for their help in flow cytometry. The FACS Aria instrument was funded by NIH grant S10-1S10RR02933801. We thank Benjamin Abrams, UCSC Life Sciences Microscopy Center, RRID: SCR_021135 for technical support during image acquisition and processing. We thank Paloma Medina for helping with bioinformatic analysis. We thank Professor Terry Rabbitts, from the Institute of Cancer Research London, for providing Lmo2-CreERT2 targeted mouse strain. We also thank the animal facility core members for animal maintenance. We thank members of the Sikandar lab for critical feedback on the manuscript. The authors declare no competing interests. This work was supported by the NIH/NCI R37CA269754 to S.S.S, NIH/NIGMS R25 PREP 5R25GM104552-09 NIH Grant T34 GM140956-01 to M.J and 5T32GM133391 to I.J.F.

## AUTHOR CONTRIBUTIONS

V.H.A and S.S.S conceived and designed the study. V.H.A and M.J. performed most of the experiments and with assistance from I.J.F under the supervision of S.S.S. V.H.A., M.J., A.O. and S.S.S. wrote the manuscript. All authors commented on the manuscript.

## COMPETING INTERESTING

The authors declare no competing interests.

---

**Supplementary Figure 1:**
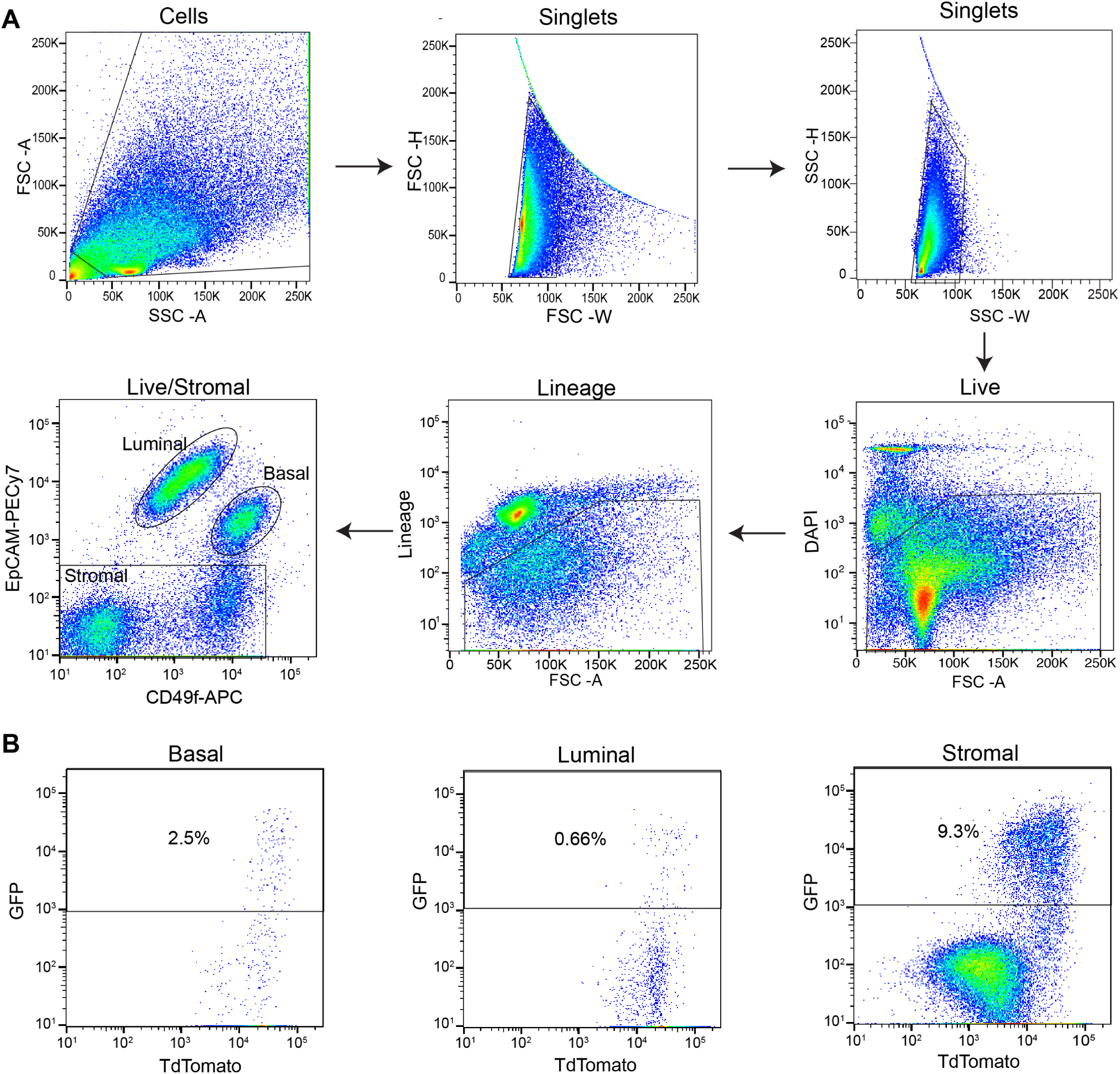
Gating strategy for lineage tracing mice *Lmo2^creERT^*/*Rosa26^mTmG^*. Mammary gland from transgenic mice *Lmo2^creERT2^*/*Rosa26^mTmG^*were digested and analyzed with flow cytometry (BD FACSAria). (**A**) Cells were isolated from debris by SSC-A x FSC-A gate and single cells were gated based on the FSC-H x FSC-W and SSC-H x SSC-W. Live cells were gated based on the live/dead cell stain DAPI. Immune cells were gated out with the lineage (lineage -) cocktail (CD45, CD31 and Ter119). Basal epithelial cells were gates based on CD49f^hi^/EPCAM^med/low^ and luminal cells gated with CD49f^low^/EPCAM^hi^. (**B**) Gates for GFP+/TdTomato+ of basal, luminal, and lineage depleted cells.

**Supplementary Figure 2:**
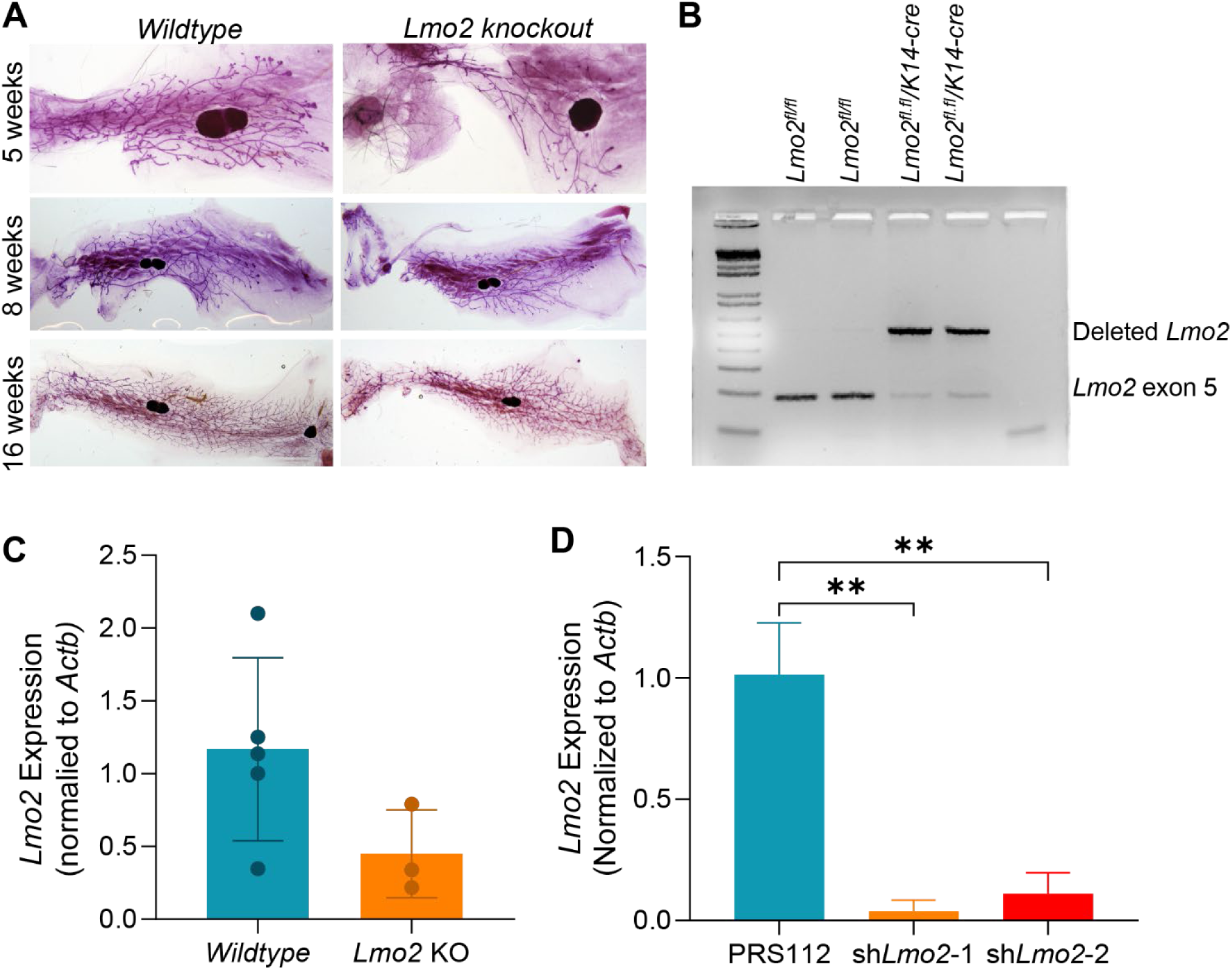
*Lmo2* knockout mice genotyping and *Lmo2* knockdown in mouse mammary epithelial cells. (**A**) Wholemount images of mammary glands from wildtype *Lmo2^fl/fl^* and *Lmo2^fl/fl^; Krt14-Cre* mice at 5, 8, and 16 weeks of age, stained with Carmine Alum. (**B**) Representative electrophoresis gel showing PCR amplification of *Lmo2* exon 5 in *Lmo2^fl/fl^* and *Lmo2^fl/lf^; Krt14-Cre mice*.(C) Quantitative PCR analysis of mammary gland tissue from *Lmo2^fl/fl^* (n=5) and *Lmo2^fl/fl^; Krt14-Cre* (n=3). (D) Quantitative PCR analysis of *Lmo2* knockdown in mouse mammary epithelial cells. Statistical significance was calculated using an unpaired t-test. * p < 0.05, ** p < 0.01, *** p < 0.001, **** p<0.0001.

**Supplementary Figure 3.**
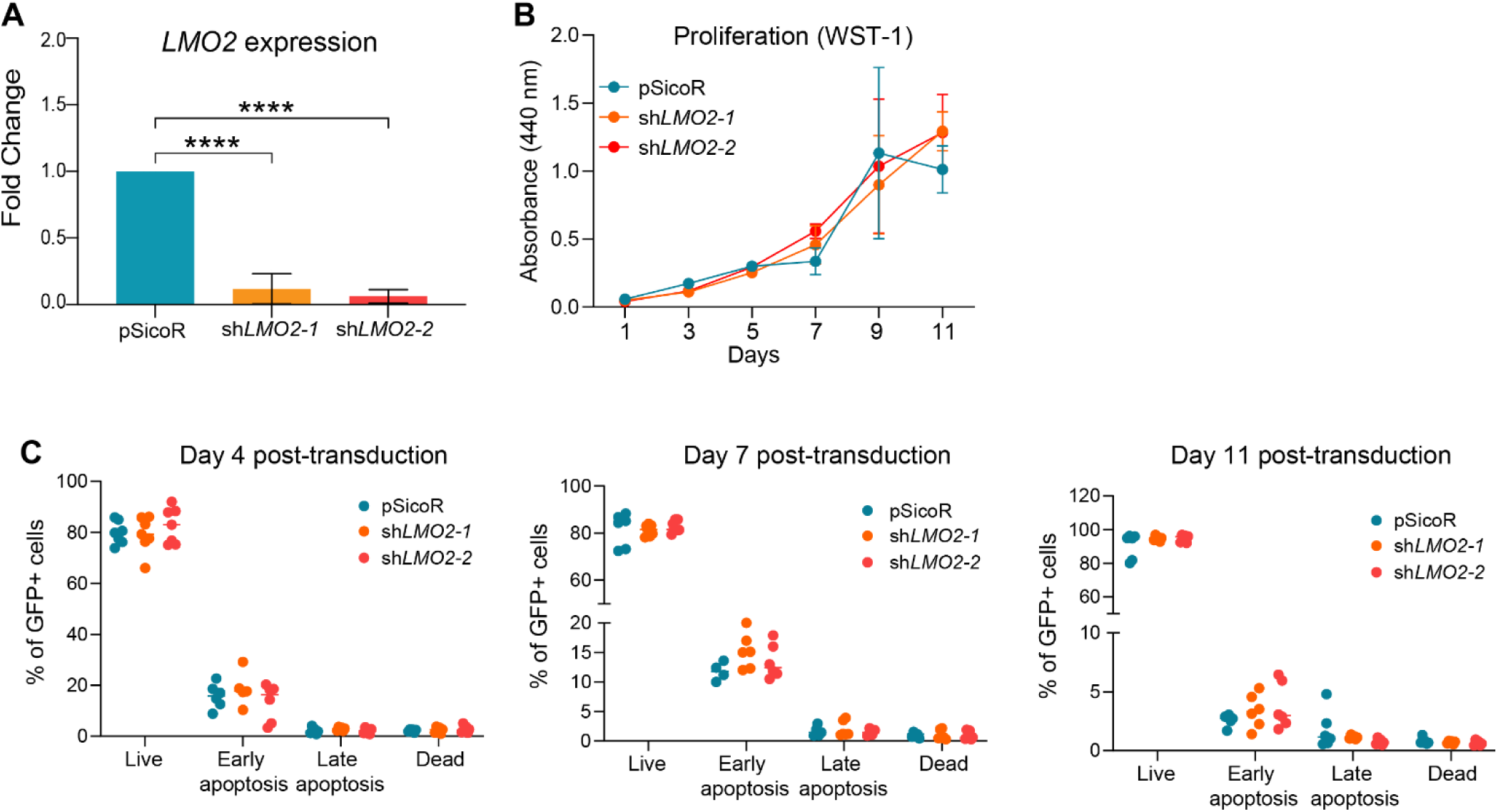
Quantification of viability and apoptotic activity in *LMO2* knockdown MCF10A cells. (**A**) Quantitative PCR confirming *LMO2* knockdown in MCF10A cells. (**B**) Cell viability assay of control and *LMO2* knockdown MCF10A cells over 11 days (n=3). (**C**) Apoptosis analysis of control and *LMO2* knockdown MCF10A cells at 4, 7, and 11 days post-transduction (n=6), showing proportions of live, early apoptotic, late apoptotic, and dead cells based on GFP+ cells. Data are shown as mean ± SD. Statistical significance was determined by using ordinary one-way ANOVA with multiple comparison test. * p < 0.05, ** p < 0.01, *** p < 0.001, **** p<0.0001.

**Supplementary Table 2.** Antibodies.

| **Antibody** | **Catalog** | **Concentration** |
| --- | --- | --- |
| DAPI | Invitrogen #D1036 | 1:10,000 |
| PE anti-human CD326 | BioLegend #324205 | 1:100 |
| APC anti-human/mouse CD49f | BioLegend #313616 | 1:200 |
| PE/Cy7 anti-human CD146 | BioLegend #361007 | 1:200 |
| PE/Cy7 anti-mouse CD326 | BioLegend #118216 | 1:100 |
| PacBlue anti-mouse CD31 | BioLegend #102422 | 1:200 |
| PacBlue anti-mouse CD45 | BioLegend #103126 | 1:200 |
| PacBlue anti-mouse TER-119 | BioLegend #116232 | 1:200 |
| APC Annexin V | BioLegend #640920 | 1:20 |

